# Proteome-wide QTL mapping enables gene-protein-phenotype metabolic network construction in a genetically diverse MASLD mouse model

**DOI:** 10.64898/2026.07.26.740836

**Authors:** Margaret Lea Robinson, Wenyu Liu, Giorgia Benegiamo, Miller T. Williams, Gordon I. Smith, Samuel Klein, Johan Auwerx, Joshua J Coon

## Abstract

The limited treatment options available for the estimated 38% of adults worldwide affected by Metabolic Dysfunction Associated Steatotic Liver Disease (MASLD) are largely due to an incomplete understanding of the complex molecular networks underlying disease pathogenesis. To dissect the genetic architecture and proteomic regulation underlying MASLD, we generated a genetically diverse mouse cohort through a four-way cross of founder strains with varying susceptibility to liver disease, producing 444 F2 mice with a spectrum of phenotypes and genotypes. Utilizing deep proteomic profiling of the livers of this population, we identified quantitative trait loci (QTL) for 2,652 proteins, spanning over 4,000 unique genomic loci, and distinguished cis- and trans-acting regulatory mechanisms. Integrating proteomic, genomic, and phenotypic data reveals key regulatory loci and candidate proteins influencing disease progression, creating a mineable proteogenomic resource for MASLD research. We utilize this resource to identify the E3 ubiquitin ligase Ubr1 as a candidate central regulator connecting proteostasis and lipid metabolism, with genetic polymorphisms that may alter its abundance and predispose to metabolic dysfunction. This study demonstrates how high-throughput, deep proteome profiling integrated with QTL mapping can reveal complex gene-protein networks governing MASLD susceptibility and progression, offering novel insights for biomarker discovery and therapeutic targeting.

## INTRODUCTION

Protein interactions and regulatory networks underlie a vast degree of phenotypic variation in cells, tissues, and organisms, including disease states. Susceptibility to such diseases can be encoded genetically, often involving a multitude of genomic regions giving rise to dysfunction.^1^ One such system is found in the genetic drivers underlying the progression of metabolic dysfunction-associated steatotic liver disease (MASLD). MASLD is a condition which causes high levels of fats to accumulate in the liver, often correlated to a combination of dietary and genetic risk factors.^2^ Metabolic dysfunction-associated steatohepatitis (MASH) is a more severe form of the disease marked by steatosis, inflammation, and liver damage.^3,4^ While these earlier stages of the disease are reversible with diet and lifestyle changes, patients are often asymptomatic and thus undiagnosed until they have progressed to cirrhosis and/or hepatocellular carcinoma (HCC), at which point the damage is no longer reversible.^5^ MASLD and its more severe forms affect around 38% of adults worldwide,³ yet the two approved therapies for MASH demonstrate incomplete efficacy, with semaglutide inducing MASH resolution in 63% of patients and resmetirom producing clinically meaningful responses in only approximately 25– 30% of treated patients.^6,7,8^ This points to an incomplete mechanistic understanding of the disease. Studies attempting to determine the degree of heritability of MASLD put estimates between 30-60%.^9^ However, better understanding of the genetic underpinnings of this disease requires study of the gene networks involved, as the metabolic pathways affected are highly complex and interconnected.^10^ With this understanding, we can then start to find targets to assess risk, provide more precise biomarkers for each stage of the disease, or even discover drugs that can reverse liver damage.

Compounding the difficulties due to the complexity of the disease, current animal models fail to encompass the spectrum of phenotypes seen in disease progression from MASLD to MASH, and fail to model many of the hallmark symptoms of MASH like hepatocyte ballooning and fibrosis.^11^ Further, these models often have a singular genetic background and rely on diet-induced phenotypes, preventing gene-phenotype association studies. Thus, to study these relationships, we have here created a four-way cross of mouse strains both susceptible and resistant to MASLD/MASH to create a cohort of 48 F0, 24 F1 and 444 F2 mice (Figure 1B).^12^ With this model encompassing large amounts of genetic and phenotypic diversity in liver metabolic disease susceptibility, we can map Quantitative Trait Loci (QTL).^13^ This forward genetic analysis can allow for determination of the genomic markers that correlate to quantitative phenotypic traits, such as body weight or steatosis scoring.^14,15^

**Figure 1.**
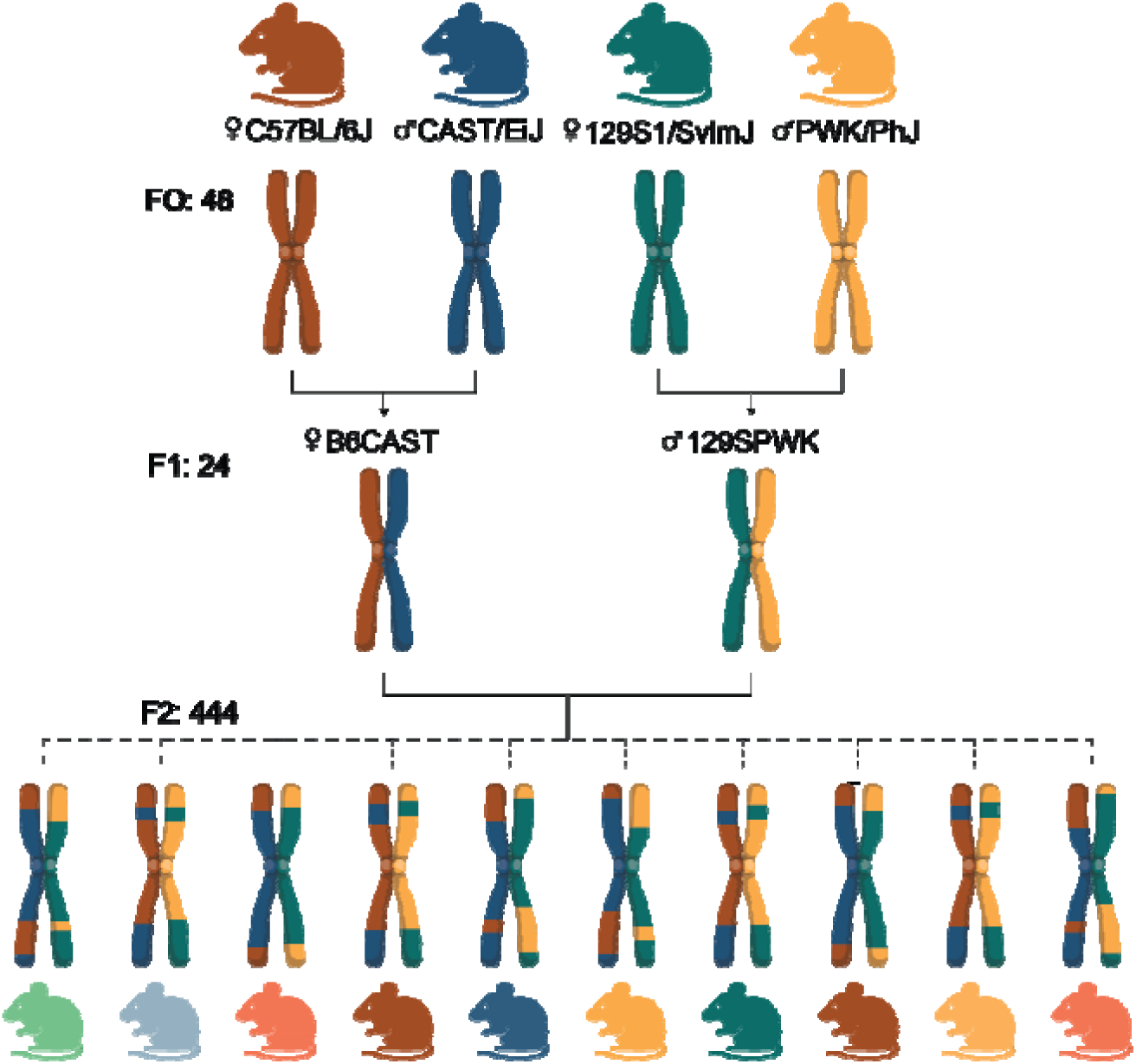
A genetically diverse four-way mouse cross. Breeding scheme for the four-way cross. C57BL/6J females were crossed with CAST/EiJ males to generate B6CAST F1 offspring, and 129S1/SvlmJ females were crossed with PWK/PhJ males to generate 129SPWK F1 offspring. B6CAST F1 females were then crossed with 129SPWK F1 males to generate the F2 generation, in which recombination produces a mosaic of all four founder genomes. The cross yielded 48 F0, 24 F1, and 444 F2 mice used in pQTL mapping, with a continuous spectrum of genotypes and liver disease phenotypes.

To create a bulk map of gene-protein relationships, we couple this approach with deep proteomics data to determine genetic factors influencing the abundance of each protein being expressed in liver tissue.^16^ This allows elucidation of genetic polymorphisms that give rise to altered protein expression in liver metabolism pathways. QTL analysis, however, relies on large sample cohorts to encompass enough genetic diversity for high resolution mapping, as well as phenotypic diversity for high statistical power.^15^ Historically, this has not been compatible with the relatively low throughput of global proteome analysis, compared to more mature technologies like transcriptomics. Studies of these magnitudes are often plagued by artifacts of analytical variation, and generally sacrifice proteome coverage and precise quantitation in exchange for increased speed of acquisition. The utility of proteomics data in QTL mapping has been well demonstrated, however, as much of proteome variation is modulated post transcriptionally.^13,17,18,19^

In the past several years, mass spectrometry (MS) instrumentation has drastically pushed the boundaries of MS scanning speed, expanding our ability to sample complete proteomes in record analysis times and minimized sample preparation.^20^ For example, the Orbitrap Astral, with a scanning speed of over 200Hz, has enabled the detection of over 10,000 proteins in just thirty minutes, approaching a near complete human proteome.^21–24^ Equally important to MS instrumentation, search algorithms for the processing of Data Independent Acquisition (DIA) MS spectra have advanced in identification, quantitation, and false discovery control, such that DIA analysis has nearly surpassed the more traditional Data Dependent Acquisition in both power and utilization.^21,25^ With this deep proteome sampling capability, we can more consistently detect and quantify many more proteins, including low abundance proteins like those involved in signaling. This can allow for higher power in determining both proximal and distal genetically encoded regulation of proteins through QTL mapping.

Here, we leverage this latest generation MS technology to quantify 12411 proteins across 444 experimental samples (9790 per sample on average), all analyzed in under one month. From this data, we mapped 2652 proteins to over 4,000 distinct genomic loci. By overlaying these pQTL with tissue and organismal level phenotype data, we begin to unravel the complexities of genetic control of the proteome in metabolically dysfunctional liver. Using deep proteomics integrated with systems genetics, we establish a proteome-wide framework for connecting genetic variation, protein abundance, and metabolic phenotypes. This resource enables identification of regulatory relationships among known MASLD-associated proteins while nominating new candidate regulators for future mechanistic investigation.

This work serves as a resource to the metabolism research community, as one of the largest proteomics studies of liver tissue metabolism to date. Combined with the genetic diversity and deep phenotyping of the mouse cohort, this pQTL dataset can be used to examine differences in many metabolic traits stemming from liver metabolism. This data is available to the public at MAPLE: MASLD Associated Proteomic Loci Explorer (https://mlr98.shinyapps.io/pQTL_MASLD/), an online interactive data viewer.

## RESULTS

### F2 generation of mice exhibit high degree of genetic, phenotypic and proteomic diversity

Of the four parental mouse strains (F0 mice) from the cross (Figure 1A), housing at thermoneutrality and feeding a high fat Western Diet was sufficient to induce liver damage in PWK/PhJ, C57BL/6J, and 129S1/SvImJ.^12^ PWK/PhJ mice were the most prone to develop MASH, and the only strain to develop fibrosis. Conversely, CAST mice were almost completely resistant to liver damage under these conditions. F2 mice, in turn, exhibited disease phenotypes all across this range seen in the parent mice.^26^

Utilizing the fast scanning speed of the Orbitrap Astral,^21^ proteome data from the liver tissue of all mice was acquired in under 45 minutes per sample with single-shot DIA LC-MS/MS, totaling just over 3 weeks of acquisition time for the 444 experimental liver samples (along with an additional 97 associated control and blank samples). An average of 9790 proteins were detected per sample (Figure 2A), and a total of 12411 proteins were detected across all samples (Figure 2B). These proteins were identified using the genomes of all four F0 mouse strains, to avoid bias towards identifications of proteins differing in any one strain. Variation in protein quantitation between different mice of the same strain was lowest in the F0 and F1 generations, and increases in the F2 generation, in correspondence with the degree of genetic diversity present across these mice (Figure 2C). 9776 of these proteins were detected in 50% of samples, and were thus used in QTL analysis.

**Figure 2.**
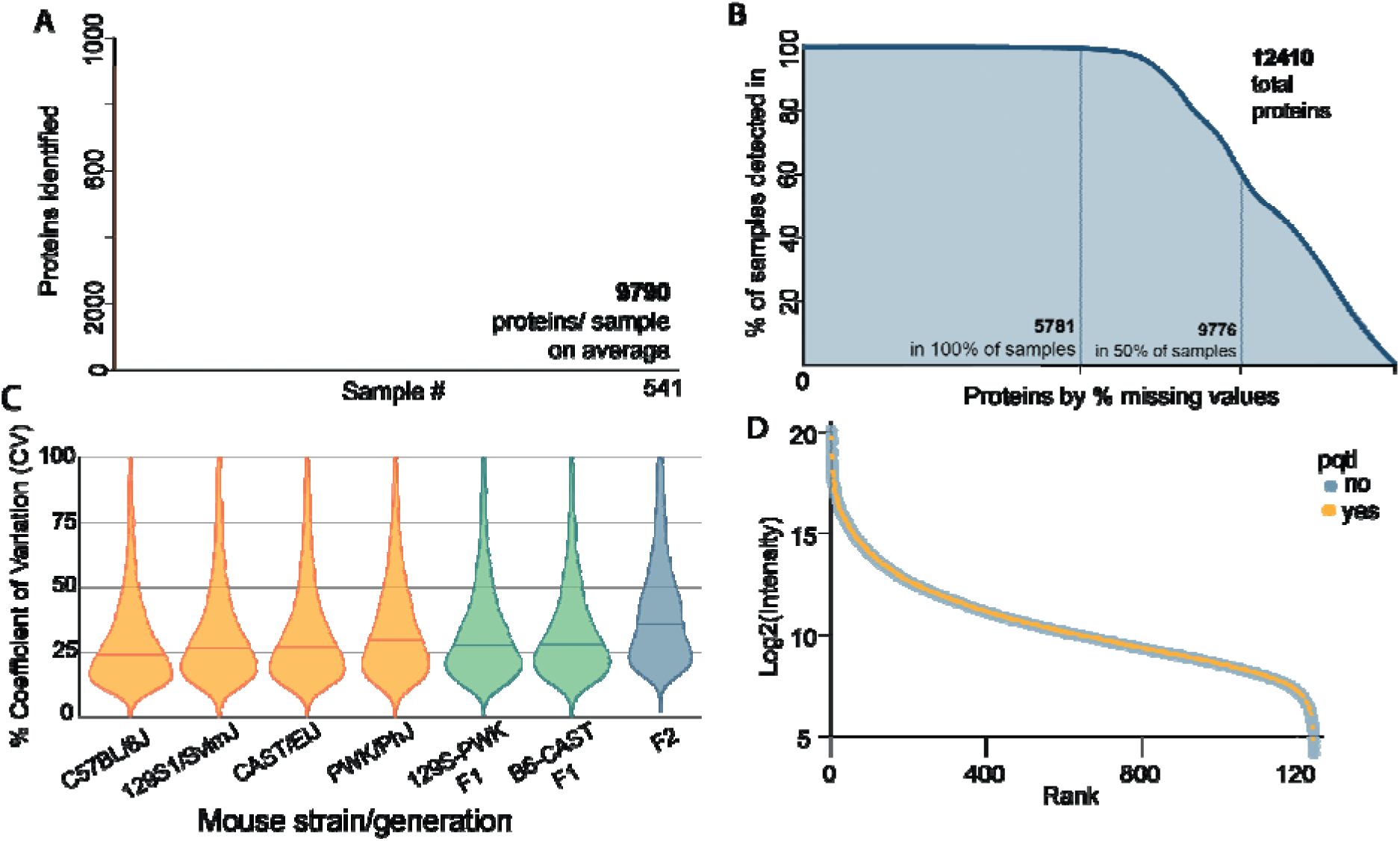
Proteomic profiling of liver tissue across the mouse cohort. (A) Number of proteins identified in each of the 541 liver samples (444 experimental, 97 control) analyzed by single-shot DIA-MS, with an average of 9,790 proteins detected per sample. (B) Cumulative distribution of the 12,410 total proteins identified across the dataset, ranked by the percentage of samples in which each was detected; 5,781 proteins were detected in 100% of samples and 9,776 in at least 50% of samples, the threshold used for downstream QTL analysis. (C) Percent coefficient of variation (CV) in protein abundance across the four F0 founder strains, the two F1 crosses, and the F2 generation. CV increases progressively with genetic diversity, from the inbred founder strains through the F1 hybrids to the genetically heterogeneous F2 population. (D) Protein abundance (log2 intensity) ranked from highest to lowest, colored by whether the protein had a detectable pQTL (yes/no), showing that pQTL were identified across the full dynamic range of the proteome, including both highly and lowly abundant proteins.

### Over 4000 loci identified to regulate protein abundances through QTL analysis

One or more pQTL were detected for 2652 of these proteins, including proteins among the most highly and lowly expressed in the proteome (Figure 2D). Over 4000 distinct loci were identified, with an average size of about 3M base pairs, equivalent to ∼30 genes (Sup. Figure 1A). Note, QTL analyses were conducted separately on the male and female populations to avoid the confounding influence of sex in the mouse cross. Of the 6638 unique pQTL found in the female dataset (7154 in male), 2418 were identified within ±1M base pairs of the gene encoding that protein, and are thus annotated as cis pQTL (2269 cis in male) (Figure 3B, Sup Figure 2B). Those 3924 outside of this range are denoted as trans, representing pQTL from regions of the genome distant from the encoding gene of a protein (4560 trans in male). 1823 pQTLs were shared between the male and female datasets (Sup Figure 3). pQTL found with high significance and agreement in both the male and female datasets include genes previously strongly associated with the progression of liver disease such as Pnpla3, Hsd17b13, and Mboat7,^27,28,29^ as well as novel loci of interest (Figure 3A, Sup Fig 2A).

**Figure 3.**
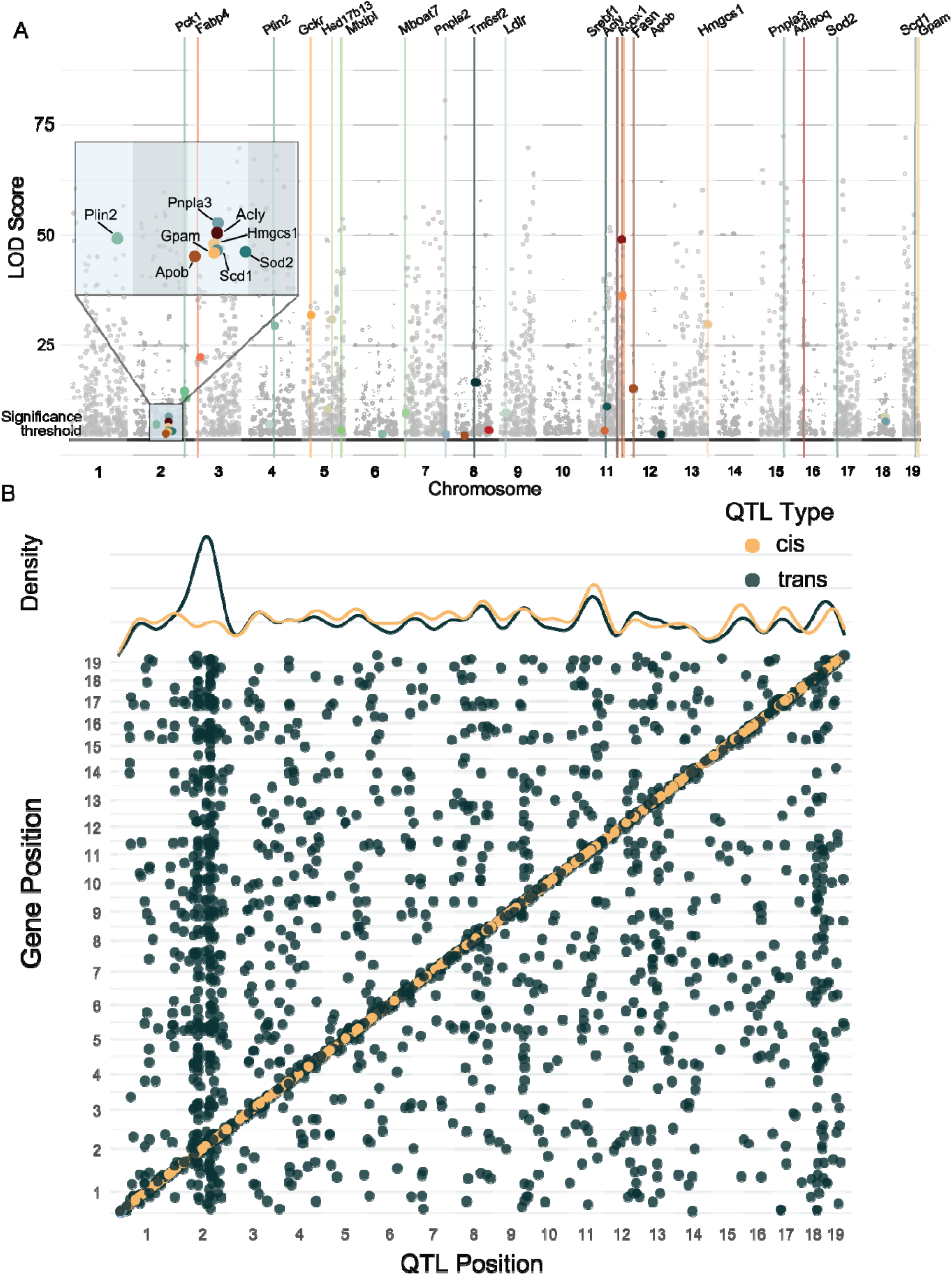
Genome-wide mapping identifies over 4,000 loci regulating liver protein abundance. (A) Manhattan plot of LOD scores for all detected protein quantitative trait loci (pQTL) across the 19 mouse autosomes in the female mouse population (identical figures from male mice are found in Sup Fig 2). Each point represents a pQTL for an individual protein; colored points highlight pQTL for proteins with established roles in lipid metabolism and MASLD (e.g., Pck1, Fabp4, Plin2, Pnpla3, Acly, Hmgcs1, Scd1, Sod2, Adipoq, among others), and vertical lines indicate their corresponding gene. Inset magnifies a cluster of these pQTL on chromosome 2. The horizontal line denotes the genome-wide significance threshold. (B) Scatter plot comparing the genomic position of each pQTL (x-axis) to the genomic position of the gene encoding the corresponding protein (y-axis), with cis pQTL (within 1 Mb of the encoding gene) in orange and trans pQTL (more distant) in teal. Marginal density plots show the distribution of cis and trans pQTL along the genome, with a pronounced trans pQTL hotspot on chromosome 2.

### Online interactive data viewer serves as resource for MASLD research

To make this resource broadly accessible to the research community, we developed an interactive, web-based data viewer MAPLE: MASLD Associated Proteomic Loci Explorer (https://mlr98.shinyapps.io/pQTL_MASLD/) allowing users to visualize the full pQTL and proteomic datasets generated in this study. The first two tabs display genome-wide pQTL maps for the female and male mouse populations, mirroring Figures 3A and 4, plotting the peak maximum LOD score for each significant pQTL with a minimum LOD threshold slider and a protein search tool; hovering over a point reveals the protein name, QTL location and type, LOD score, and tau correlation to steatosis (the male dataset, corresponding to Sup. Figure 2, is shown on the second tab). A third tab, corresponding to Figure 5A, provides a scatterplot tool for the mouse proteomics dataset, allowing users to plot any combination of measured phenotypes and protein abundances against one another, colored by sex. A fourth tab, mirroring Figure 7, extends this functionality to the human liver cohort, enabling users to plot protein abundance against liver disease diagnosis (MASLD, MASH, or other metabolic disease), colored by sex. A final tab, corresponding to Sup. Figure 5, directly compares cross-species disease relevance for all pQTL-mapped proteins by plotting human tau correlation to steatosis against mouse tau correlation, with point size reflecting LOD score and color denoting QTL type, with protein search and minimum LOD filters available. By enabling researchers to explore pQTL relationships, co-regulation patterns, and cross-species concordance for any protein of interest we hope this tool will extend the impact of this dataset well beyond the present study and serve as a lasting resource for the broader metabolism research community.

**Figure 4.**
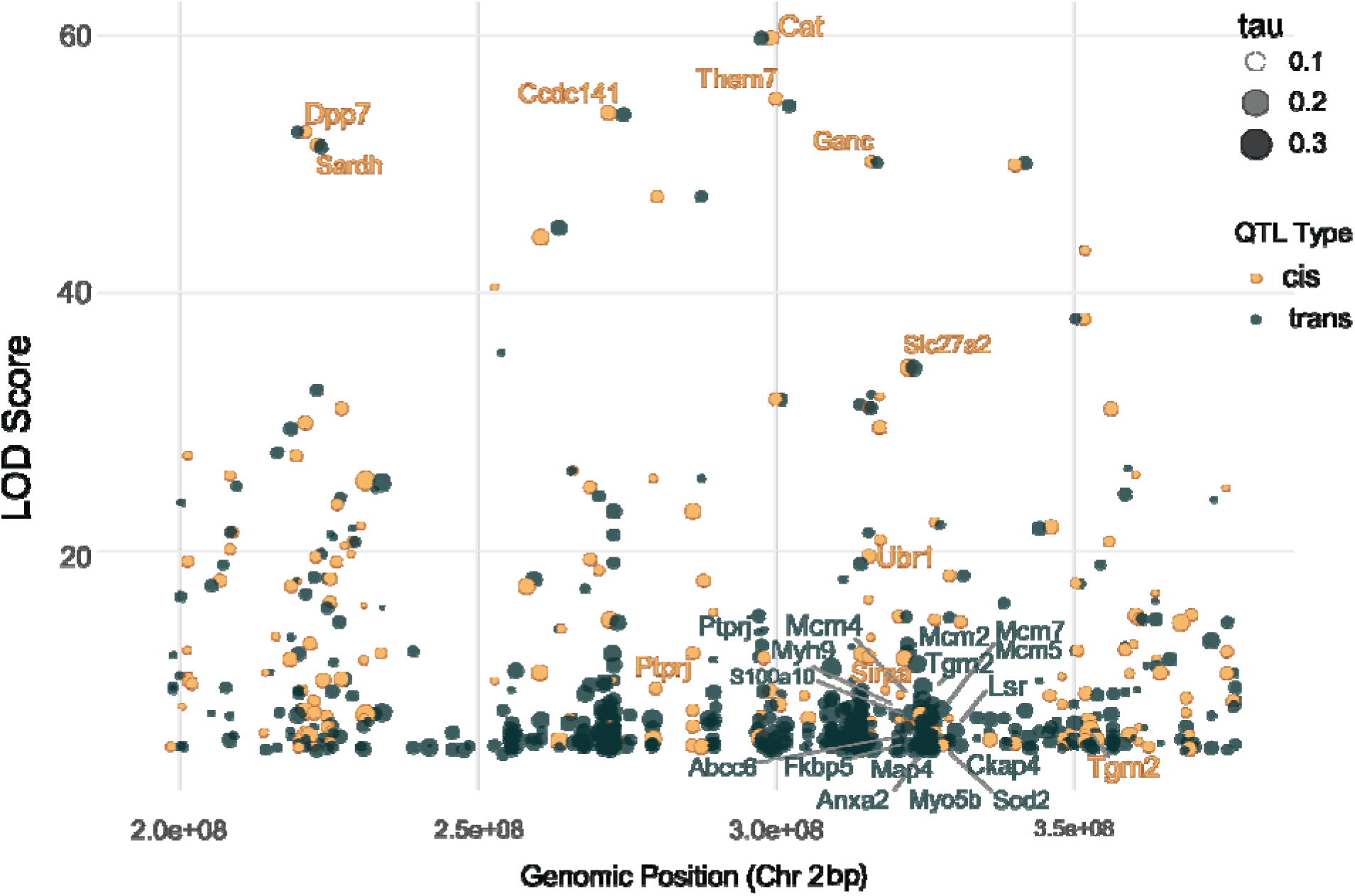
A trans pQTL hotspot on chromosome 2 is associated with steatosis and lipid metabolism genes. LOD score plotted against genomic position along chromosome 2 for all detected pQTL, with cis pQTL shown in yellow and trans pQTL in green. Point size reflects tau, the strength of correlation between protein abundance and steatosis score. Labeled genes include high-LOD cis pQTL (e.g., Dpp7, Sardh, Ccdc141, Cat, Them7, Ganc, Slc27a2, Ubr1) and trans-regulated genes clustering near the Ubr1/steatosis locus around 124 Mb, including members of the MCM helicase complex (Mcm2, Mcm4, Mcm5, Mcm7) and cytoskeletal and proteostasis-associated proteins (Sirpa, Tgm2, Ptprj, Myh9, S100a10, Fkbp5, Anxa2, Sod2, among others).

**Figure 5.**
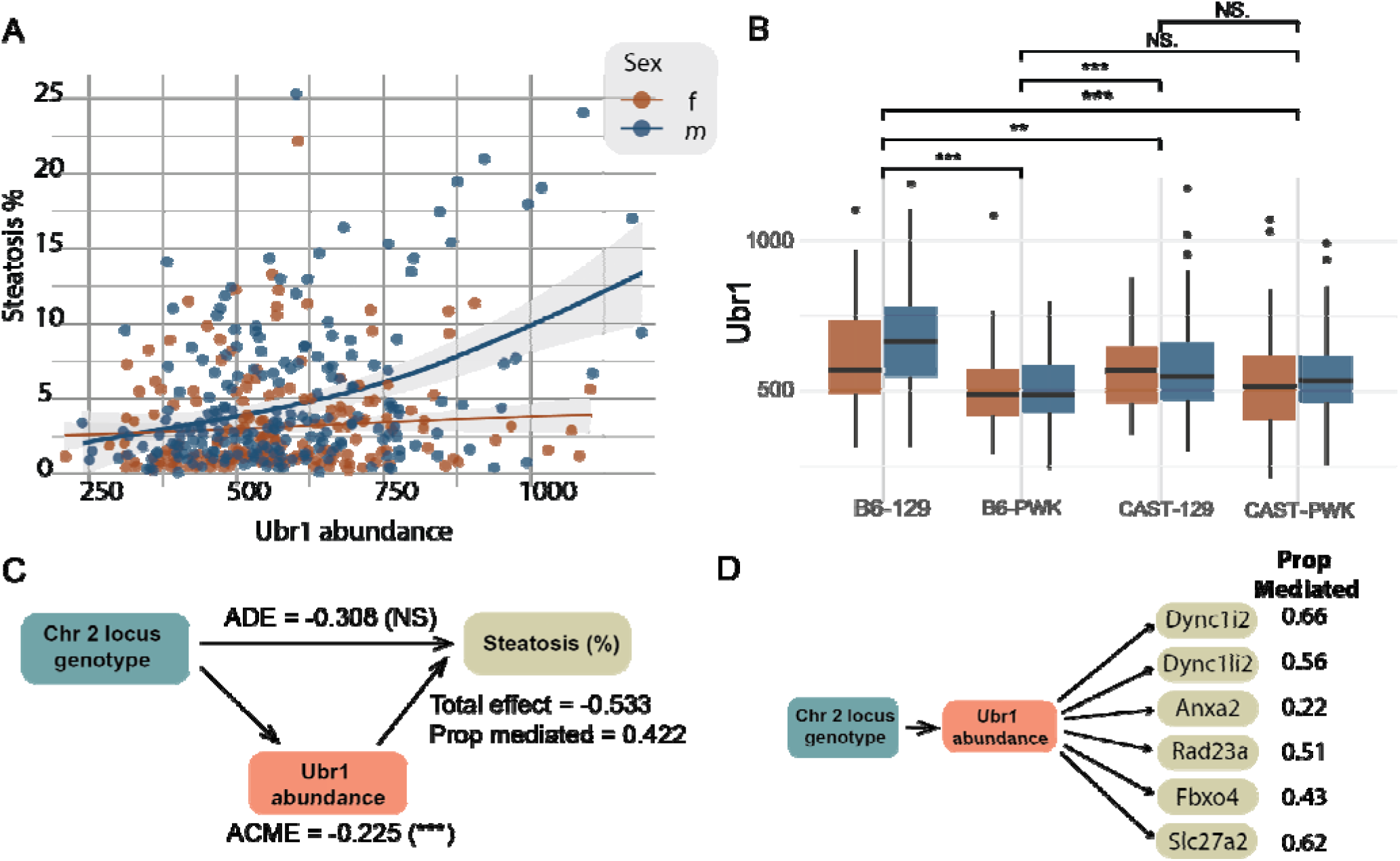
Ubr1 abundance mediates the chromosome 2 locus effect on steatosis. (A) Ubr1 protein abundance vs. histological steatosis score across F2 mice, by sex (brown, female; blue, male). The positive association is stronger in males. Shading, 95% CI. (B) Ubr1 abundance by chromosome 2 parental allele combination (B6-129, B6-PWK, CAST-129, CAST-PWK) and sex. The 129 allele is associated with higher Ubr1 abundance than PWK, regardless of the paired founder (**p < 0.01, ***p < 0.001; NS, not significant). (C) Mediation analysis of Ubr1 abundance on the chromosome 2 locus–steatosis relationship. ADE = −0.308 (NS); ACME (via Ubr1) = −0.225 (***p < 0.001); total effect = −0.533; proportion mediated = 0.422. (D) Proportion of the locus’s effect on candidate Ubr1 substrates mediated through Ubr1 abundance, ranging from 0.22 (Anxa2) to 0.66 (Dync1i2), supporting Ubr1 as a functional link between this locus and downstream proteostasis targets.

### Integration of liver phenotyping with pQTL identifies overlapping genomic hotspots for steatosis and protein abundances

Utilizing this resource, we investigated potential causal genes and proteins driving steatosis susceptibility. Because pQTL analysis is conducted using only proteomic and genomic information and is therefore agnostic to disease phenotypes, we incorporated histopathological measures of liver disease to identify genetic drivers specifically associated with metabolic dysfunction. Specifically, we calculated Kendall’s tau correlations between protein abundance and histological steatosis scores, which quantify hepatic lipid accumulation, for individual mice while accounting for sex as a covariate. Many proteins show significant association with steatosis, 1681 of which also have a detected pQTL. A large proportion of these pQTL are detected in trans from the protein’s gene, and are found on chromosome 2 (Figure 4).

We see many microtubule-associated proteins with high LOD pQTL correlating to steatosis scoring. Microtubule stability is well known to play an important role in liver function, like in the case of Wilson’s disease, where their destabilization positively correlates to the development of fibrosis.^30,31^ Additionally noteworthy among these steatosis-associated proteins are several members of the MCM complex (Mcm2-5, Mcm7), which are upregulated in HCC and are associated with poor prognosis^32^. This complex forms a helicase important to DNA replication, as well as damage sensing and repair, and suppresses cellular senescence to facilitate HCC progression^33^. All proteins of the complex excluding Mcm6 correlate positively to steatosis and share a trans pQTL locus on chromosome 2. Importantly, the mice in this cross have not yet progressed to HCC and many have not yet even developed fibrosis, indicating this complex may have a role very early in the disease progression.

During phenotypic QTL analysis, steatosis scoring showed two high LOD peaks on chromosome 2 at 68 Mbp and 124 Mbp.^26^ These same regions harbored a large clustering of trans pQTL including trans regulation of Pnpla3, (Figure 3AB) mutations of which are known drivers of MASLD^34^. Thus, we have examined this region of the genome closer to identify possible underlying molecular changes caused by the genes encoded here. To identify which genes within the QTL peaks may be driving the steatosis phenotype, we first examined the cis pQTL in the regions.

### Resource-guided prioritization identifies UBR1 as a candidate mediator of steatosis susceptibility

Under the steatosis QTL peak at 124Mb of chromosome 2, we detected 30 cis pQTL (Figure 4). To determine if any of these proteins are likely to be causal for this steatosis QTL, we employed mediation analysis between the abundance of these proteins and steatosis scoring. Of the genes in this region with significant mediation scores, only the protein Ubr1 also had a cis pQTL. Particularly emphasized in the male mice, the expression of Ubr1 is positively correlated to steatosis scoring (Figure 5AB) and it is mediating around 42% of the effect of this locus on steatosis variability (Figure 5C). This, while a high degree of mediation, is also reflective of a likely multi-factor locus, which we would expect to see with high levels of trans regulation of many proteins^35^. Ubr1 is an E3 ubiquitin ligase, shown to directly sense and bind dietary branched chain amino acids and flag Plin2 for degradation^36^. Under amino acid deprivation, Ubr1 is inactivated and Plin2 levels are elevated, causing higher levels of lipid accumulation in the cell^37^. Interestingly, we see a positive correlation between Plin2 and Ubr1 protein levels (Figure 6A), indicating a potentially more complex regulatory relationship or an overall downstream increase in Plin2 as a result of increased steatosis.

**Figure 6.**
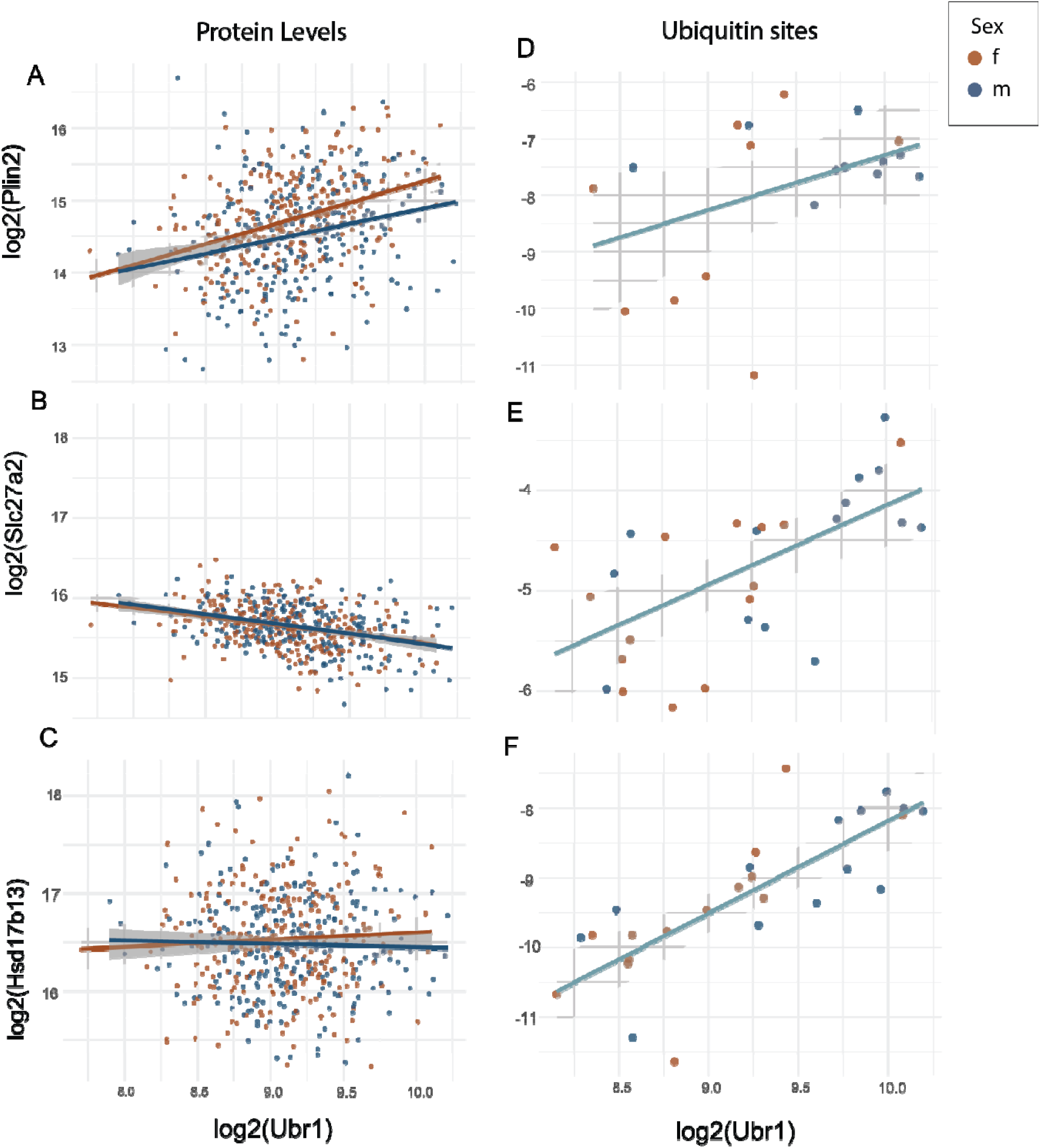
Ubr1 abundance correlates with both protein levels and ubiquitination of downstream lipid metabolism targets. Relationship between log2(Ubr1) protein abundance (x-axis) and either total protein level (A-C, left) or relative abundance of a specific ubiquitination site (D-F, right) for four lipid metabolism-associated proteins, colored by sex (brown, female; blue, male). (A, D) Plin2 protein level and ubiquitination site, both increasing with Ubr1 abundance. (B, E) Slc27a2 protein level, which decreases with Ubr1 abundance, and its ubiquitination site, which increases with Ubr1 abundance. (C, F) Hsd17b13 protein level and ubiquitination site. In each case, ubiquitination site abundance increases with Ubr1 levels even where total protein abundance is flat or inversely related, consistent with Ubr1-mediated ubiquitination of these targets independent of their overall expression.

**Figure 7.**
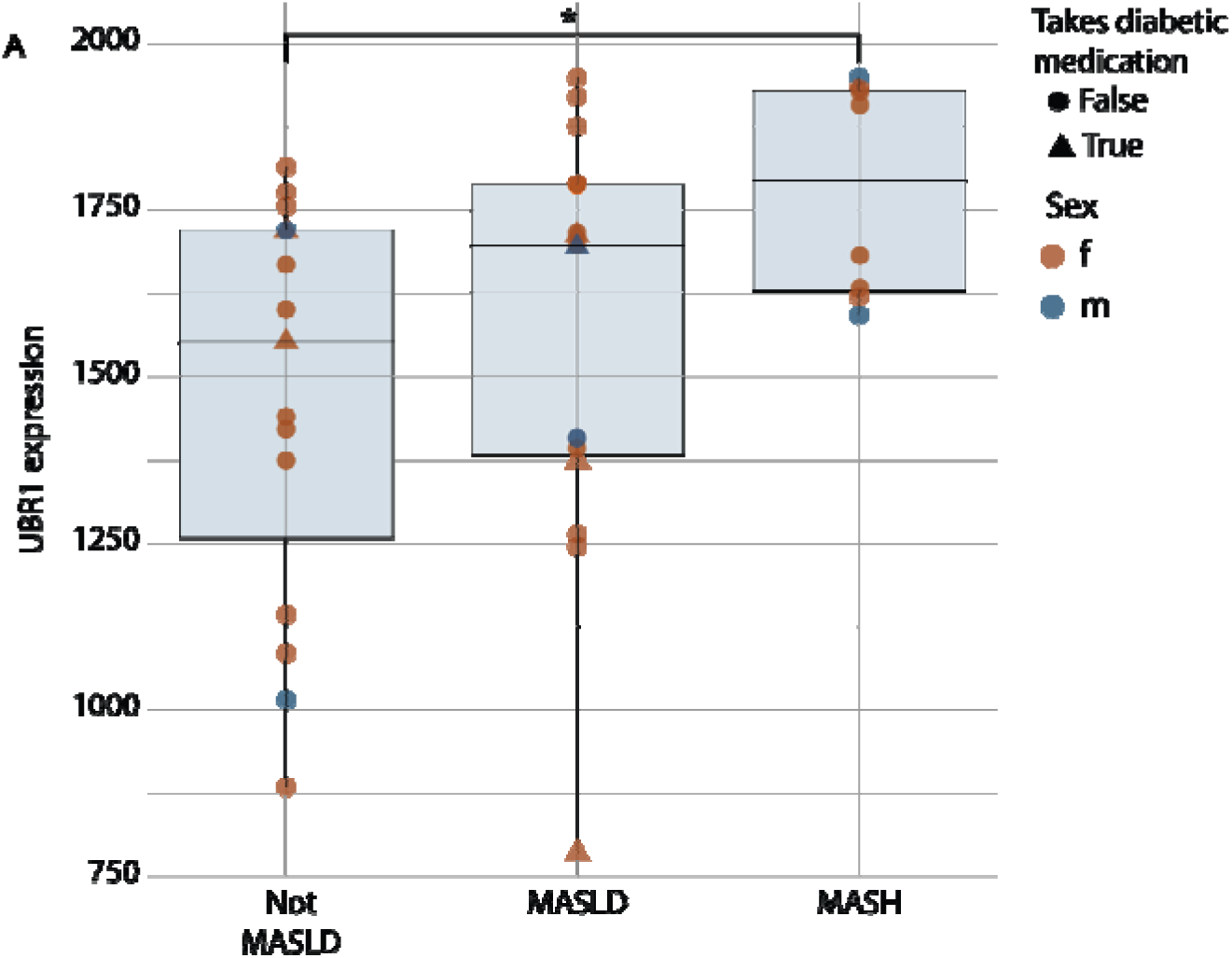
UBR1 expression is elevated in human MASLD and MASH and correlates with steatosis severity. UBR1 expression in human liver biopsies stratified by diagnosis: not MASLD, MASLD, and MASH. UBR1 expression increases with disease severity, with the highest levels observed in subjects with MASH. Points are colored by sex (female, brown; male, blue) and shaped by whether the subject was taking diabetic medication (circle, no; triangle, yes). UBR1 expression increases with steatosis grade.

Because we observed this region of the genome to also harbor many trans pQTL, we conducted additional mediation analysis to determine whether variation in Ubr1 abundance mediated the effects of the locus on distal protein abundances^35^. Indeed, Ubr1 significantly mediated the abundance of numerous trans-regulated proteins spanning diverse biological processes, including intracellular trafficking, cytoskeletal organization, ubiquitin-dependent proteostasis, and lipid metabolism (Figure 5D). Among the most strongly mediated proteins were multiple components of the cytoplasmic dynein motor complex, cytoskeletal regulators, and proteins involved in ubiquitin-dependent protein turnover, suggesting that genetic variation at the Ubr1 locus broadly remodels cellular transport and protein homeostasis networks. ^38–40^.

Additionally, several metabolic proteins involved in fatty acid activation and sterol biosynthesis were significantly mediated, indicating that UBR1-associated proteome remodeling extends to pathways directly involved in hepatic lipid metabolism. Among the proteins most negatively correlated to both Ubr1 and steatosis is Slc27a2, a very long chain fatty acid (VLCFA) transporter implicated in steatotic liver disease, expressed primarily on the peroxisome.^41,42^ Slc27a2 expression shows an inverse relationship with steatosis score and Ubr1 abundance (Figure 6B), likely causing decreased VLCFA transport into peroxisomes for beta oxidation. The resulting accumulation of VLCFA inside of cells and in lipid droplets is a hallmark of MASLD. Together, these findings support UBR1 as a central regulator linking genetic variation to coordinated changes in intracellular transport, protein quality control, and metabolic function.

### Ubiquitin site enrichment identifies direct targets of Ubr1

To directly probe the pathways through which Ubr1 may be propagating its effect on steatosis, we conducted ubiquitin site profiling using di-glycine remnant enrichment.^43^ This was done to identify proteins Ubr1 may be modifying for degradation or other regulation. We selected 30 samples for enrichment, including a variety of high and low Ubr1 expressors, as well as high and low steatosis scorers. Ubr1 is primarily known to participate in the proteasomal degradation system, following the N-end rule in which it ubiquitinates N-termini with destabilizing residues ^44–46^, thus playing a role in general unspecific proteome maintenance for misfolded proteins.

Across the 30 samples, we detected at least one ubiquitin site on 3204 distinct protein groups, 2367 of which have multiple sites detected. Strikingly, UBR1-associated ubiquitination sites were highly enriched in proteins involved in both proteostasis and in metabolic pathways, including ribosomal proteins, proteasome subunits, and chaperonins, as well as key regulators of lipid metabolism (Figure 6, Sup Fig 4). Several of these proteins exhibited multiple ubiquitination sites with consistent directional effects, suggesting coordinated regulation at the protein level and lack of site specificity for ubiquitin tagging in many cases. Multiple sites on Plin2 increase with abundance alongside increasing Ubr1, even when normalized for protein level abundance (Figure 6A,D). This is evidence that Ubr1 is likely fulfilling its canonical role of targeting Plin2 for degradation, but Plin2 abundance is more strongly determined by alternative factors like increased expression. Nine of the proteins with correlated Ub sites also had detected pQTL within the region of the Ubr1 and steatosis peaks. This included enzymes and transporters such as Scd1, Acsl5, Slc27a2, and the lipid droplet-associated protein Hsd17b13 (Figure 6D-F, Sup Fig 4), further evidence that Ubr1 expression is varying the abundance of these key proteins involved in lipid handling in the liver. These data suggest Ubr1 may have an additional, more specific role as a central regulator in lipid metabolism.

### Human MASH livers show increased expression of UBR1

To validate findings from our mouse cohort and test relevance to human health, we collected proteomics data on a cohort of human livers using the same methodology. Similar to the mouse cohort, these clinical samples presented a range of MASLD symptoms, including some subjects with minimal or no hepatic steatosis. Across the 38 livers, we detected 10,149 unique protein groups, with an average of 9848 detected per sample. We again calculated Kendall’s tau correlation coefficients to determine the proteins that may be involved in MASLD pathogenesis. Despite lower sample numbers and high heterogeneity in the human cohort, we detected many proteins that correlated similarly between mice and humans. 52% of orthologs showed sign concordance of tau between mice and humans, indicating the same directionality of their relationship to steatosis (Sup Fig 5).

Notably, UBR1 again showed moderate (Kendall’s tau = 0.185) correlation to liver fat, and is significantly upregulated in subjects with MASH diagnosis (Figure 7). Additionally, we find that UBR1 abundance is associated with coordinated changes in proteostasis-related pathways in human liver (Figure 8A). Proteins involved in ubiquitination, including HUWE1, UBR4, ARIH1, and HECTD3, as well as deubiquitinases such as USP9X and USP24, showed positive correlations with UBR1. In contrast, molecular chaperones and co-chaperones, including BAG1 and HSPE1, were negatively correlated. We also observed inverse correlations with central metabolic enzymes, including the glycolytic enzyme PGK1 and the TCA cycle component DLST. While limited by cohort size and clinical heterogeneity, these data suggest that UBR1 abundance in human liver is associated with a coordinated proteomic state characterized by increased ubiquitin-mediated protein turnover and altered metabolic activity (Figure 8B).

**Figure 8.**
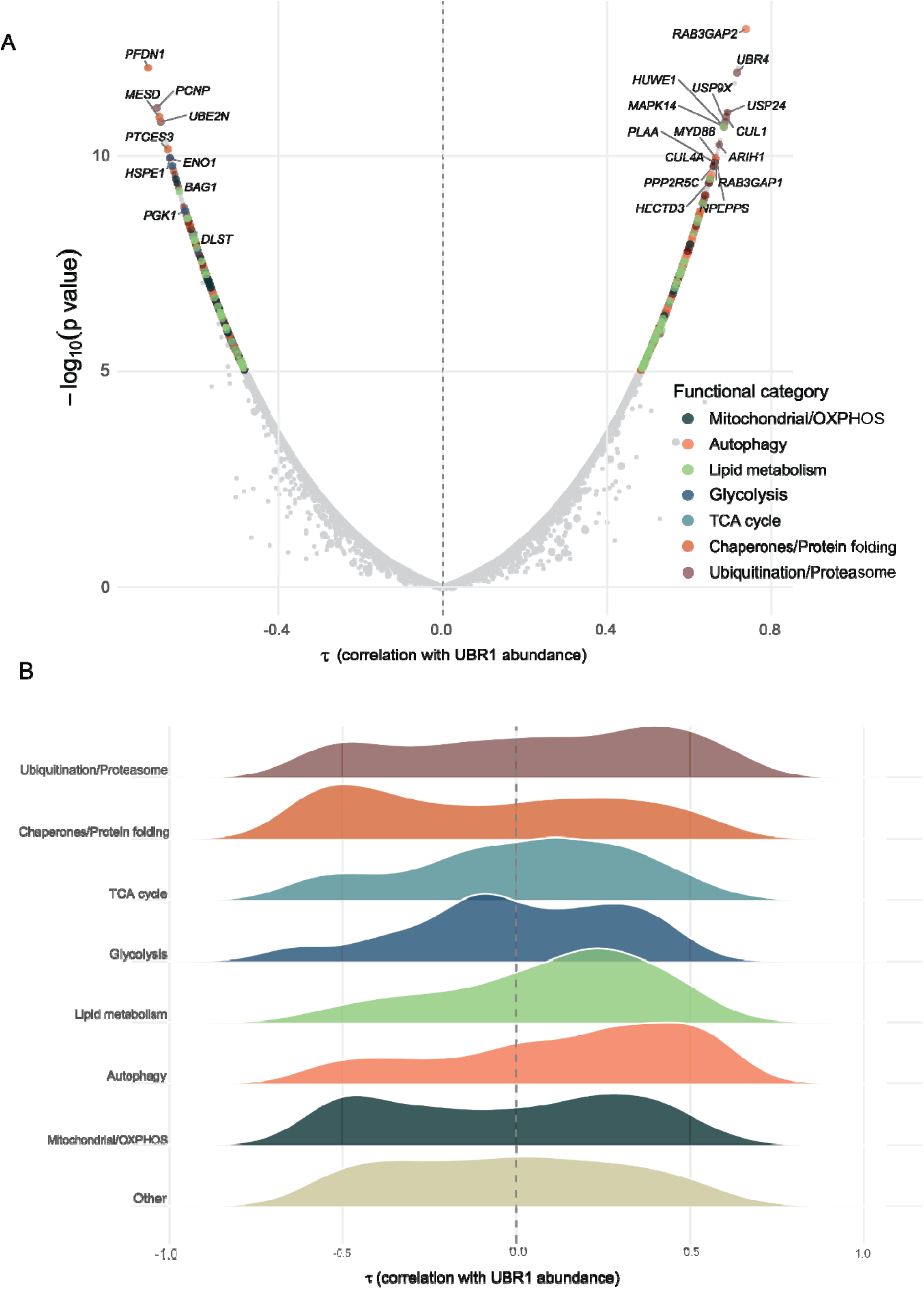
UBR1 abundance is associated with coordinated changes in proteostasis and metabolic pathways in human liver. (A) Volcano plot of Kendall’s tau correlation between protein abundance and UBR1 abundance (x-axis) against -log10(p-value) (y-axis) across the human liver proteome. Proteins are colored by functional category (mitochondrial/OXPHOS, autophagy, lipid metabolism, glycolysis, TCA cycle, chaperones/protein folding, ubiquitination/proteasome). Proteins positively correlated with UBR1 include ubiquitin ligases and deubiquitinases (e.g., UBR4, HUWE1, USP9X, USP24, CUL1, ARIH1, RAB3GAP1/2), while negatively correlated proteins include molecular chaperones (e.g., HSPE1, BAG1) and glycolytic/TCA cycle enzymes (e.g., PGK1, ENO1, DLST). (B) Ridgeline plot showing the distribution of tau values for proteins in each functional category, illustrating that ubiquitination/proteasome and autophagy proteins skew toward positive correlation with UBR1, while chaperone/protein folding proteins skew negative.

Search of the whole proteome data for di-glycine remnant on ubiquitin modified peptides detected 177 sites, further revealing that ubiquitination sites most strongly associated with UBR1 abundance in human liver span a diverse set of proteins involved in proteostasis, intracellular trafficking, and metabolism (Figure 9, Sup Fig 6). Highly correlated sites were identified on proteins such as ACOX1, SLC2A2, and ADH1C, indicating that UBR1-associated ubiquitination extends to key metabolic enzymes. In addition, ubiquitination of proteins involved in vesicle trafficking and membrane dynamics, including AP1B1 and NAPA, was positively associated with UBR1 levels, suggesting a potential role in regulating protein localization and secretion. Consistent with its established function, UBR1 was also associated with ubiquitination of canonical proteostasis factors such as HSP90AB1. Together, these results indicate that UBR1 abundance in human liver is linked to a coordinated ubiquitination program spanning protein turnover, trafficking, and metabolism, consistent with and extending observations from the mouse model.

**Figure 9.**
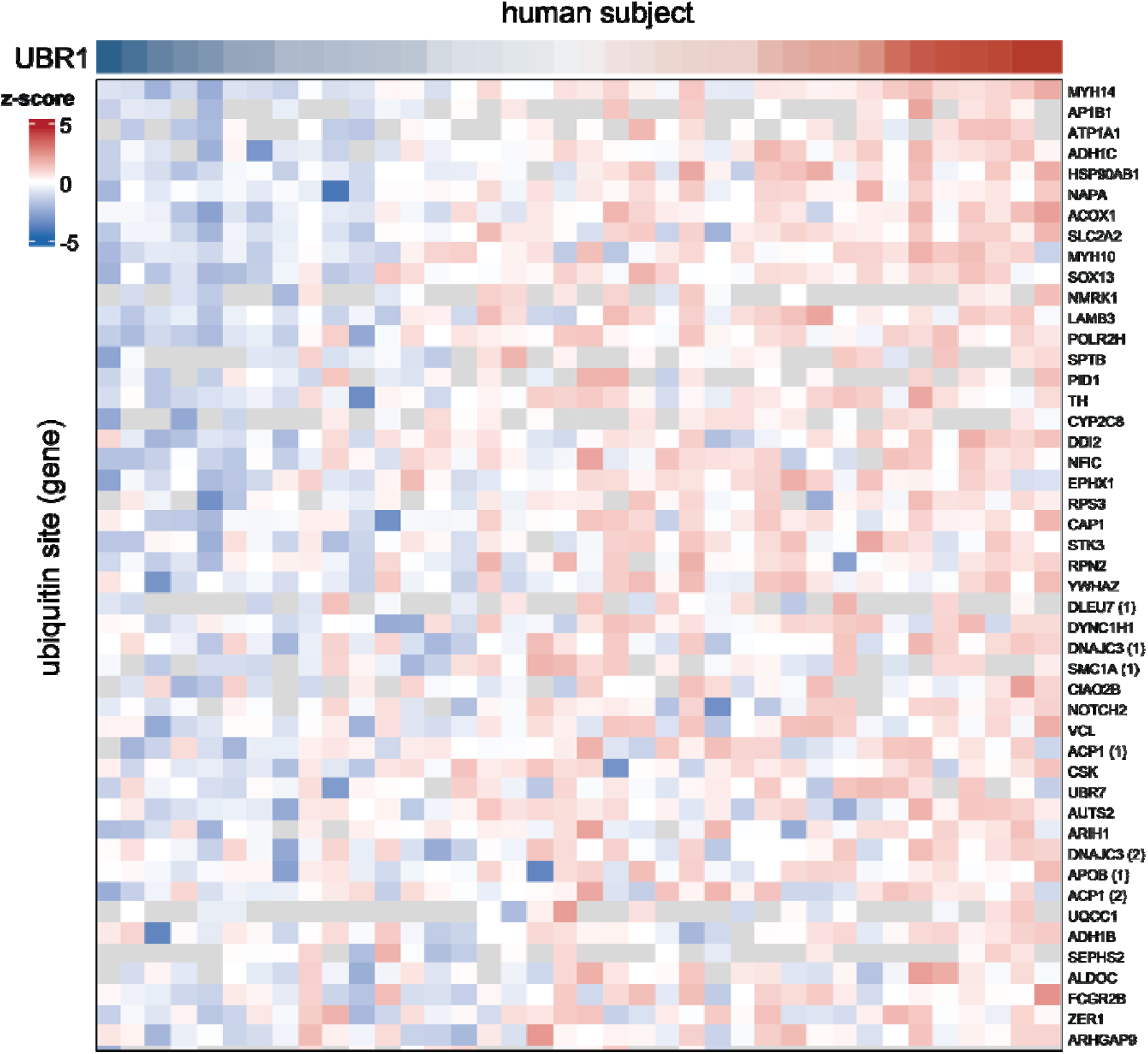
UBR1 abundance correlates with a coordinated ubiquitination signature across human liver subjects. Heatmap of top 50 most correlated ubiquitination sites to UBR1 abundance (row z-score) across human liver subjects (columns), ordered left to right by increasing UBR1 abundance (color bar, top). Rows correspond to individual ubiquitination sites, labeled by gene name, with numbers in parentheses distinguishing multiple sites on the same protein. Sites shift from lower (blue) to higher (red) relative abundance with increasing UBR1, spanning proteins involved in proteostasis (e.g., HSP90AB1, UBR7, ARIH1), metabolism (e.g., ACOX1, SLC2A2, ADH1C, CYP2C8), and intracellular trafficking (e.g., AP1B1, NAPA, MYH14, MYH10).

## DISCUSSION

The complex dysregulation of metabolism underlying the progression of MASLD remains to be elucidated in full. Here, using pQTL mapping in a genetically diverse cohort of mice, we present a means to connect distinct genetic nodes of protein regulation between metabolic pathways. This can serve to elucidate the molecular manifestations underpinning tissue level phenotypes like steatosis and fibrosis, as well as the genetic contributions controlling them.

Proteins like Pnpla3 and Mboat7 correlate to the susceptibility, development, and progression of MASLD, but the underlying connections between many of these known markers are not understood. At a shared trans pQTL on chromosome 2, we see evidence that mutations in the gene Ubr1 may directly control the abundance of Scd1, Slc27a2, and Hsd17b13, among other key lipid metabolism proteins. The loss of function in these specific pathways resulting from increased ubiquitination can lead to significant bottlenecks in the storage and metabolism of free fatty acid species.^42,47,48^ From this, the accumulation of lipotoxic species in hepatocytes triggers inflammation and eventually ballooning and fibrosis, marking the progression to MASH and then cirrhosis.^49–51^ This pQTL integration approach serves to highlight such potential novel central nodes of liver metabolism, like Ubr1.

We demonstrate the utility of pQTL mapping as a tool for hypothesis generation in complex molecular networks and heritable diseases. Increases in the depth and speed of MS proteomics data acquisition further enable the connection of distal genetic regulation of protein abundances through trans pQTL by consistently detecting nearly 10,000 distinct proteins over hundreds of samples. This scale of coverage places proteomics on competitive footing with genomic and transcriptomic approaches for mapping the genetic architecture of complex disease, while offering a distinct advantage: proteins are the functional effectors of most cellular processes and sit closer to the phenotypes we ultimately care about. Because mRNA levels are often poor predictors of the corresponding protein abundance, particularly for the post-translationally regulated and rapidly turned-over proteins central to metabolic flux, deep proteomic profiling can reveal regulatory relationships that transcriptomic studies alone would miss. Such gene-protein correlations have the potential to transform diagnosis and precision treatment for MASLD and many other heritable diseases. With this approach and additional validation to follow, we can begin to elucidate novel drug targets and biomarkers, opening up new treatment possibilities for patients in the future.

A central strength of this approach lies in its use of a genetically diverse mouse population rather than a single inbred background or a genetically engineered mouse model. Studies performed in one genetic background are, in effect, studies of a single genotype, and findings from such models have frequently failed to generalize, a limitation that has fueled longstanding skepticism about the translational value of mouse work in metabolic disease. Genetic reference populations circumvent this problem by capturing the kind of allelic and phenotypic variation seen across human populations, allowing associations to be tested for robustness across many genetic contexts. When adequately powered, such populations can nominate plausible candidate genes and pathways for human metabolic disease with a degree of generalizability that single-background studies cannot offer. Our identification of Ubr1 illustrates this directly: a candidate nominated from trans pQTL mapping across genetically diverse mouse livers was independently supported by our human liver cohort, where Ubr1 abundance correlated with liver fat and ubiquitination was detected on orthologous substrate proteins. This cross-species concordance is, in itself, evidence for the value of the genetic reference population approach; it demonstrates that a mechanism identified in mice using this strategy can translate to human disease biology.

Serving as a resource for the metabolism research community, we have made our data available through the R-shiny based interactive visualization tool MAPLE: MASLD Associated Proteomic Loci Explorer (https://mlr98.shinyapps.io/pQTL_MASLD/). Without requiring any computational expertise, researchers can directly query this dataset for their own proteins of interest, examine co-regulation patterns across the genome, and assess cross-species concordance with human liver disease. We anticipate this tool will be especially valuable for groups studying lipid metabolism, proteostasis, and other pathways relevant to MASLD and broader metabolic disease, who may not otherwise have access to genetically diverse proteomic datasets of this scale, and we hope it will extend the impact of this resource well beyond the specific findings presented here.

## METHODS

### Sample Preparation

10 mg of cryopulverized liver tissue was dissolved in 3:1:1 nButanol/ Methanol/ water and sonicated at 10C for 5 minutes. Samples were centrifuged at 15000g for 5 minutes at 4C to precipitate proteins, and supernatant was removed. Protein pellet was then dissolved in 8M urea/ 100mM Tris/ 10 mM TCEP/ 40 mM CAA, and sonicated for another 5 minutes. LysC was added at 50:1 protein: protease and incubated for 4 hours at room temperature. Sample was then diluted 1:4 in 100mM Tris. Trypsin was added at 50:1 protein: protease and incubated overnight at room temperature. Samples were acidified with formic acid to pH 2 then desalted using Waters SepPak 50mg tC18 cartridges. Peptides were then dried by speedvac and reconstituted in 0.2% FA, quantified by Nanodrop (A 205), and brought to 1mg/ml.

### K-GG enrichment

K-GG motif enrichment utilized the PTMScan HS Ubiquitin/SUMO Remnant Motif (Cell Signaling Technology, Kit #59322). Enrichment protocol from the manufacturer was followed, beginning with 500ug of mouse liver peptides. Search parameters in Spectronaut mirrored those from Kumar et al 2026^52^, using DirectDIA and a library extension containing data from triplicate K-GG enrichments of 5 mg, 3 mg and 1 mg of mouse liver peptides.

### Spectral Library Generation

Peptides from all 4 F0 parent strains of mice were pooled in equal ratios. Peptides were then fractionated (Agilent 1260 Infinity BioInert LC, Waters XBridge, Peptide BEH C18, 3.5 µm, 130 Å, 4.6 mm × 150 mm column) to 16 fractions and concatenated to 8 total fractions for analysis. LC/MS methods described above were used to collect this data.

### LCMS

500ng of peptide was injected using Vanquish Neo with a 75 micron ID column with 1.5 micron C18 particles.^53^ A 30 minute separation ramped from 100% mobile phase A (0.2% Formic Acid) to 55% mobile phase B (80% acetonitrile/ 0.2% FA/ 19.8% water) before washing and equilibration. MS/MS spectra were acquired using an Orbitrap Astral with data independent acquisition (DIA). MS1 data was collected with the Orbitrap at a resolution of 240,000 from 380-980 m/z. MS2 data was collected in the Astral mass analyzer using 4m/z DIA windows with 1m/z overlap over a scan range of 380-980 m/z, and 500% AGC with 3ms max inject time.

### Proteomics Data Analysis

DIA proteomics data was analyzed using Spectronaut (v18.1) with default settings. The reference proteomes of all four F0 strains of mice were used for database searching and library generation. Proteins detected in 50% or more of samples were used in QTL analysis. All statistical analyses were conducted in R. Documentation can be found on GitHub: https://github.com/mlr121/pQTL-analysis-in-MASLD.

### Mouse husbandry and dietary regimen

Founder animals were sourced from The Jackson Laboratory and housed at the École Polytechnique Fédérale de Lausanne (EPFL) for a minimum of two generations prior to crossing. Cages held 2–5 mice under a standard 12-hour light/dark cycle, with food and water available without restriction. Starting at 6 weeks of age, animals were transferred to thermoneutral housing (30°C) in Memmert climate cabinets; one week later, at 7 weeks of age, they were switched onto a Western-style diet (Research Diets D12079B: 40% kcal from fat, 17% kcal from protein, 43% kcal from carbohydrate). Body weights were tracked every two weeks from 6 weeks of age through the end of the study.

### Necropsy and sample collection

At 24 weeks of age, after 17 weeks of Western diet exposure, animals were fasted for 4 hours in the morning and euthanized in the afternoon (between approximately 1:30 and 4:00 p.m.). Following isoflurane anesthesia, whole blood was withdrawn from the vena cava, after which animals were perfused with chilled PBS. Liver, kidney, heart, and spleen were harvested and snap-frozen in liquid nitrogen immediately upon collection; a subset of liver tissue was instead preserved in formalin or OCT compound for histological work. Collected blood was transferred to EDTA tubes and spun at 1500 × g for 10 minutes at 4°C, and the resulting plasma was snap-frozen for later use.

### RNA isolation and library preparation

Frozen liver tissue was first ground in liquid nitrogen, and approximately 10 mg of the resulting powder was disrupted in TRIzol reagent using steel beads in a TissueLyser II (2 minutes at 30 Hz). RNA was then purified with a Direct-zol-96 kit (Zymo Research). Every sample met quality thresholds for both purity (NanoDrop) and integrity (Fragment Analyzer) prior to library construction. Strand-specific libraries were generated with the SMARTER mRNA-Seq Library Prep Kit and sequenced on a DNBSEQ instrument, targeting a minimum of 50 million paired-end 100 bp reads per library. Post-sequencing FastQC reports showed no quality issues, so no read trimming was applied. Reads were mapped to the GRCm38 (C57BL/6J) reference using STAR, and transcript abundance was estimated with RSEM to produce both expected-count and CPM expression matrices; genes were retained for further analysis only if CPM exceeded 1 in at least half of all samples.

### Variant detection from RNA-seq data

Variant calling followed GATK’s RNA-seq short variant discovery pipeline (GATK v4.3.0.0). Aligned BAM files were processed through MergeBamAlignment and MarkDuplicates, then SplitNCigarReads and base quality score recalibration, before variants were called with HaplotypeCaller in its RNA-seq configuration. Hard filtering was then applied per GATK guidelines — SNPs were excluded for QD < 2.0, FS > 60.0, SOR > 3.0, MQ < 40.0, MQRankSum < −12.5, or ReadPosRankSum < −8.0, and indels for QD < 2.0, FS > 200.0, or ReadPosRankSum < −20.0 — followed by a second filtering pass in VCFtools (v0.1.16) requiring minimum depth of 3, maximum depth of 100, minimum quality of 20, minimum genotype quality of 20, no more than 50% missingness, and minor allele frequency ≥ 0.01. The variants surviving both filters formed the basis for genotype map construction.

### Building F2 genotype maps

A custom, window-based pipeline converted the filtered variant calls into F2 genotype maps. Per-sample genotypes were first normalized into matrices, with any non-standard calls marked as missing. To identify SNPs informative for distinguishing founders, a consensus genotype was computed for each of the four founder groups (C57BL/6J, 129S1/SvImJ, CAST/EiJ, PWK/PhJ); a site was kept only if every founder group’s consensus was homozygous and internally consistent. These candidate sites were then cross-checked against the two F1 crosses (129S1/SvImJ × PWK/PhJ and C57BL/6J × CAST/EiJ), requiring the F1 consensus genotype at each site to match what would be predicted from the corresponding founder pair. Each surviving SNP could then be assigned to one of four possible F2 genotype classes based on its founder consensus alleles, termed B6-129 (C57BL/6J-129S1/SvImJ), B6-PWK (C57BL/6J-PWK/PhJ), CAST-129 (CAST/EiJ-129S1/SvImJ), and CAST-PWK (CAST/EiJ-PWK/PhJ). Every F2 animal’s genotype at that site was then compared against these four expected classes: a match to exactly one class marked the site as unambiguous for that individual, a match to more than one class marked it ambiguous, and no match marked it missing.

Sliding genomic windows were then built around clusters of unambiguous SNPs. Within each window, support for each of the four genotype classes was tallied across all informative sites — unambiguous SNPs cast a full vote for their assigned class, while ambiguous SNPs split their vote evenly across all classes they were compatible with. Whichever class accumulated the most support became the window’s provisional genotype call, but this call was only accepted if it agreed with the unambiguous-SNP evidence and passed minimum thresholds for SNP count and internal consistency. Adjacent or nearby windows sharing the same accepted genotype were then merged into larger genotype blocks, provided the gap between them fell below a set distance cutoff; windows with conflicting genotypes blocked any such merging across that interval.

Any genomic stretch left unresolved by this process, including gaps flanked by blocks of the same genotype, gaps flanked by blocks of different genotypes, and the unblocked regions at chromosome ends, was sent through the same window-scoring procedure a second time, with compatible results folded into the existing blocks. For the handful of intervals that remained ambiguous, genotype assignment relied on the surrounding block context where this gave a clear answer; intervals lacking a consistent resolution, or those conflicting with neighboring evidence, were ultimately marked missing.

Finally, the curated set of genotype blocks was mapped back down to individual SNP positions to produce the finished F2 genotype map: SNP-level calls that were missing or ambiguous, but fell inside a supported block, took on that block’s genotype, whereas calls that conflicted with their block, or fell outside any supported block, were set to missing. The complete codebase and parameter choices for this pipeline are deposited in the project’s GitHub repository.

### Human Subject Enrollment

Thirty-eight people with obesity (age 44 ± 1 years) who were scheduled for Roux-en-Y gastric bypass or sleeve gastrectomy surgery at Barnes-Jewish Hospital in St Louis, MO participated in this study, which was approved by the Institutional Review Board of Washington University School of Medicine in St. Louis, MO and registered in ClinicalTrials.gov (NCT03701828). All participants provided written informed consent before enrollment and completed a comprehensive medical evaluation that included a history and physical examination and standard blood tests to determine eligibility. Potential participants were excluded if they: i) were taking any medication that could affect IHTG content; ii) had liver disease other than MASLD; iii) consumed excessive amounts of alcohol (≥3 drinks/day for men and ≥2 drinks/day for women); and/or iv) regularly used tobacco products.

### Liver biopsy and histology

Liver tissue was obtained under direct visualization at the onset of the laparoscopic bariatric surgery procedure. Samples were rinsed in 0.9% ice-cold saline with a portion of tissue fixed in 10% formalin and subsequently embedded in optimal cutting temperature (OCT) compound for hepatic steatosis, fibrosis, inflammation and hepatocellular ballooning grading according to the Nonalcoholic Steatohepatitis Clinical Research Network (NASH CRN) scoring system.^54^ The remaining tissue was frozen immediately in liquid nitrogen and stored at -80°C until processing for proteomics analysis as described above.

## ACKNOWLEDGMENTS

Funding

This work was supported by a grant from the US National Institutes of Health (NIH P41 GM108538 to JJC), the UW-Madison Biotechnology Training Program (NIH 5 T32 GM135066 to MLR), the École Polytechnique Fédérale de Lausanne (EPFL to J.A.), the European Research Council (ERC-AdG-787702 to J.A.), the Swiss National Science Foundation (31003A-179435 and 310030-214826 to J.A.), the Swiss Cancer League (KLS-5906-08-2023 to JA and GB), NIH P30 DK56341 (Washington University Nutrition Obesity Research Center) and support from the Foundation for Barnes-Jewish Hospital.

## Author contributions

Conceptualization: MLR, GB, JJC, JA; Methodology: MLR, WL, GB, GIS; Investigation: MLR, WL, GB, MTW, GIS; Visualization: MLR; Funding acquisition: JA, JJC, SK; Writing – original draft: MLR; Writing – review & editing: all authors.

## Data and materials availability

Proteomics raw data are available at MassIVE: MSV000096193. The data and metadata are searchable on https://www.systems-genetics.org/, https://mlr98.shinyapps.io/pQTL_MASLD/

