## Supplemental Figures for "Proteome-wide QTL mapping enables gene-protein-phenotype metabolic network construction in a genetically diverse MASLD mouse model"

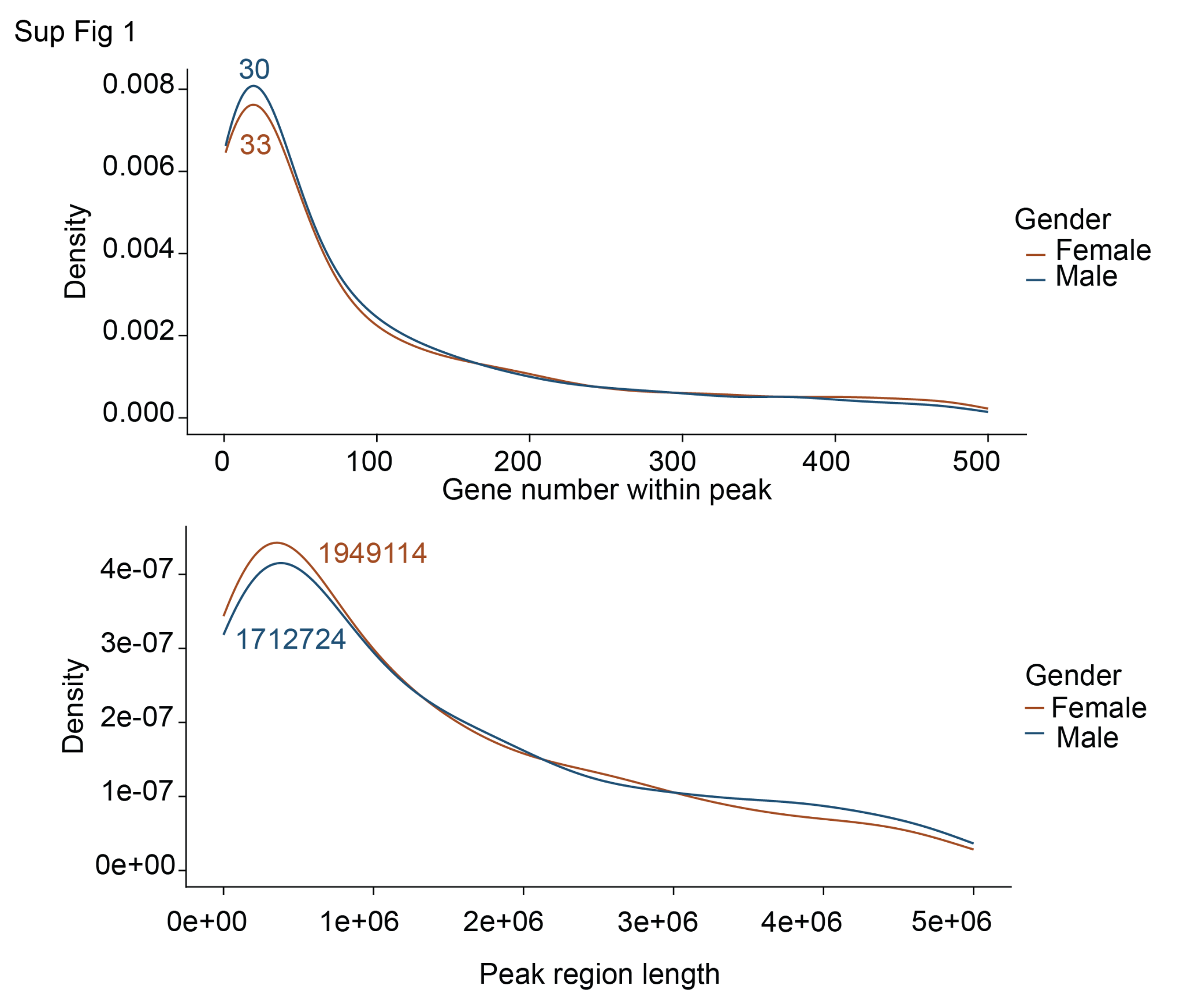

**Sup. Figure 1. pQTL peak size.**

(A) Distribution of the number of genes contained within each pQTL peak region, stratified by sex (female, brown; male, blue), with a median around 30 genes per peak. (B) Distribution of pQTL peak region lengths in base pairs, stratified by sex, with a median peak length of approximately 1.7-1.9 Mb.

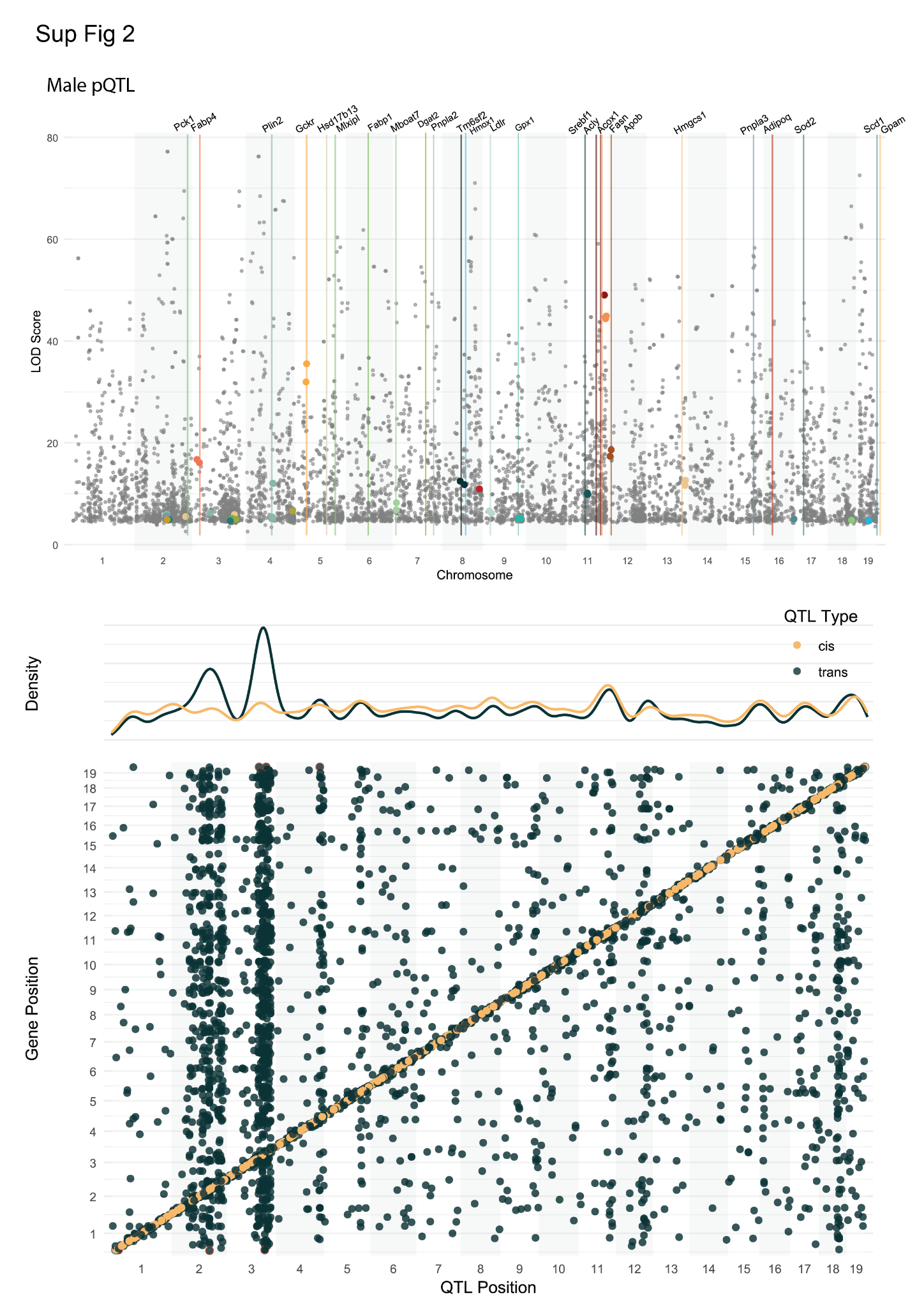
**Sup. Figure 2. Male pQTL mapping recapitulates the major genomic hotspots identified in the female analysis.**

(A) Manhattan plot of LOD scores for pQTL detected in the male-only F2 dataset, with colored points and vertical lines marking pQTL for the same lipid metabolism and MASLD-associated genes highlighted in Figure 3A. (B) Scatter plot of pQTL genomic position versus the genomic position of the corresponding gene in males, with cis pQTL in orange and trans pQTL in teal, and marginal density plots showing pQTL distribution along the genome. A prominent trans pQTL hotspot is again observed on chromosome 2, with additional peak on chromosome 3.

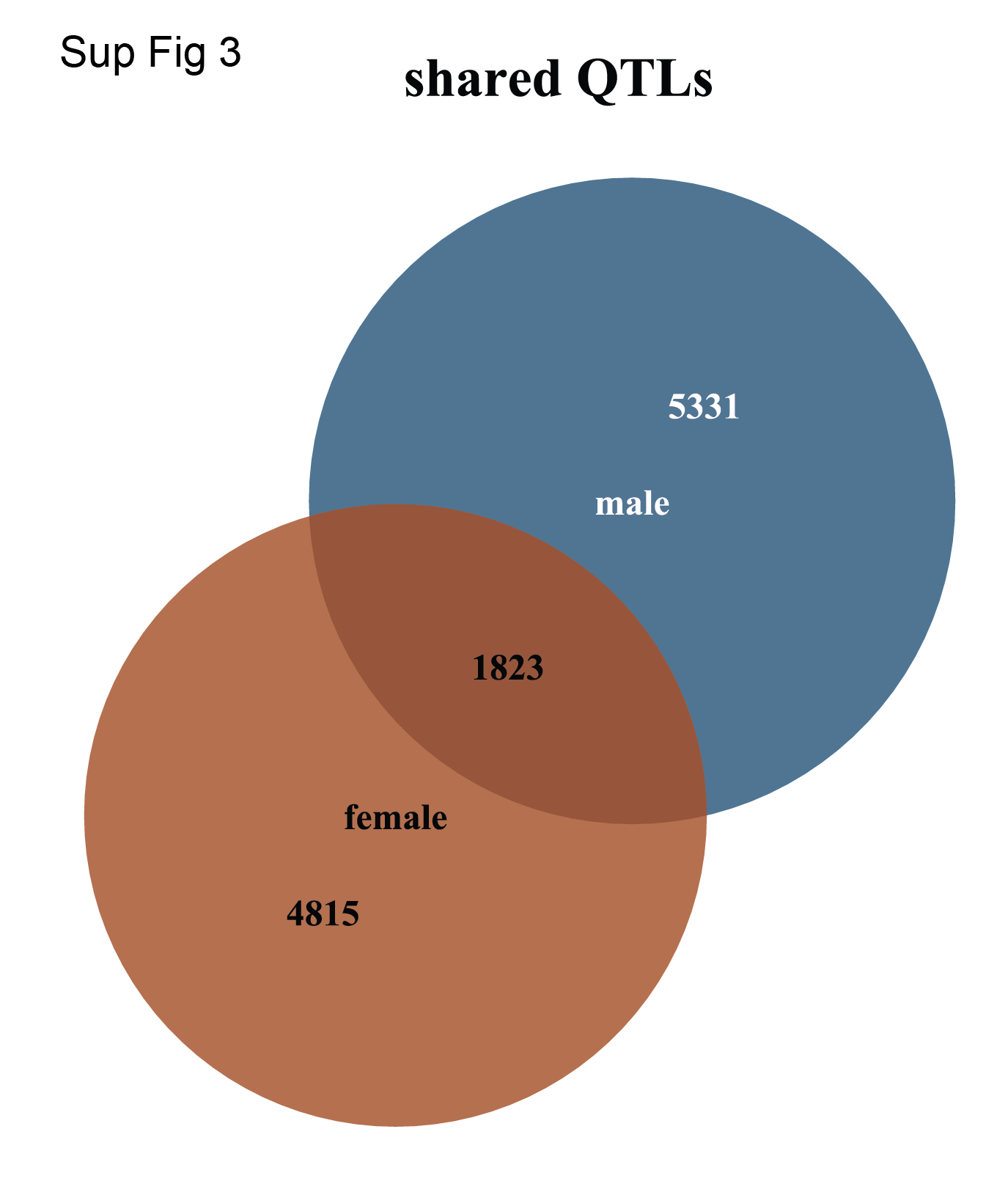

**Sup. Figure 3. Overlap of pQTL identified in male and female F2 mice.**

Venn diagram showing the number of pQTL unique to the male dataset (5,331), unique to the female dataset (4,815), and shared between both sexes (1,823).

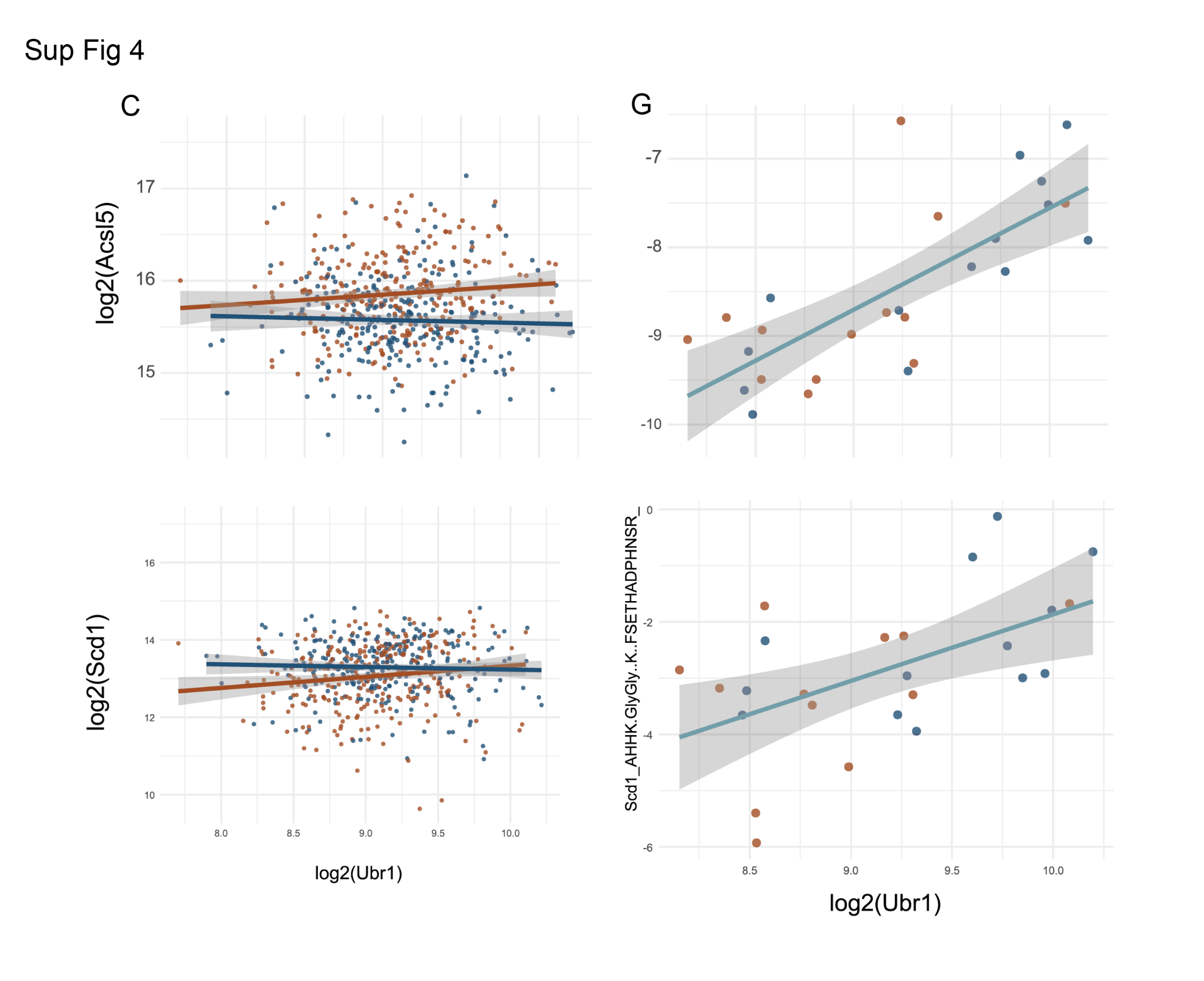

**Sup. Figure 4. Ubr1 abundance correlation with Acsl5, Scd1 protein levels and ubiquitination sites.**

(A, B) Acsl5 protein level and ubiquitination site. (C) Log2(Scd1) protein abundance plotted against log2(Ubr1) abundance across F2 mice, colored by sex (female, orange; male, blue), showing little to no overall relationship between total Scd1 protein level and Ubr1 abundance. (D) Abundance of a specific Scd1 ubiquitination site plotted against Ubr1 abundance, showing a positive correlation, indicating that Ubr1 levels are associated with increased ubiquitination of Scd1 independent of changes in total Scd1 protein abundance.

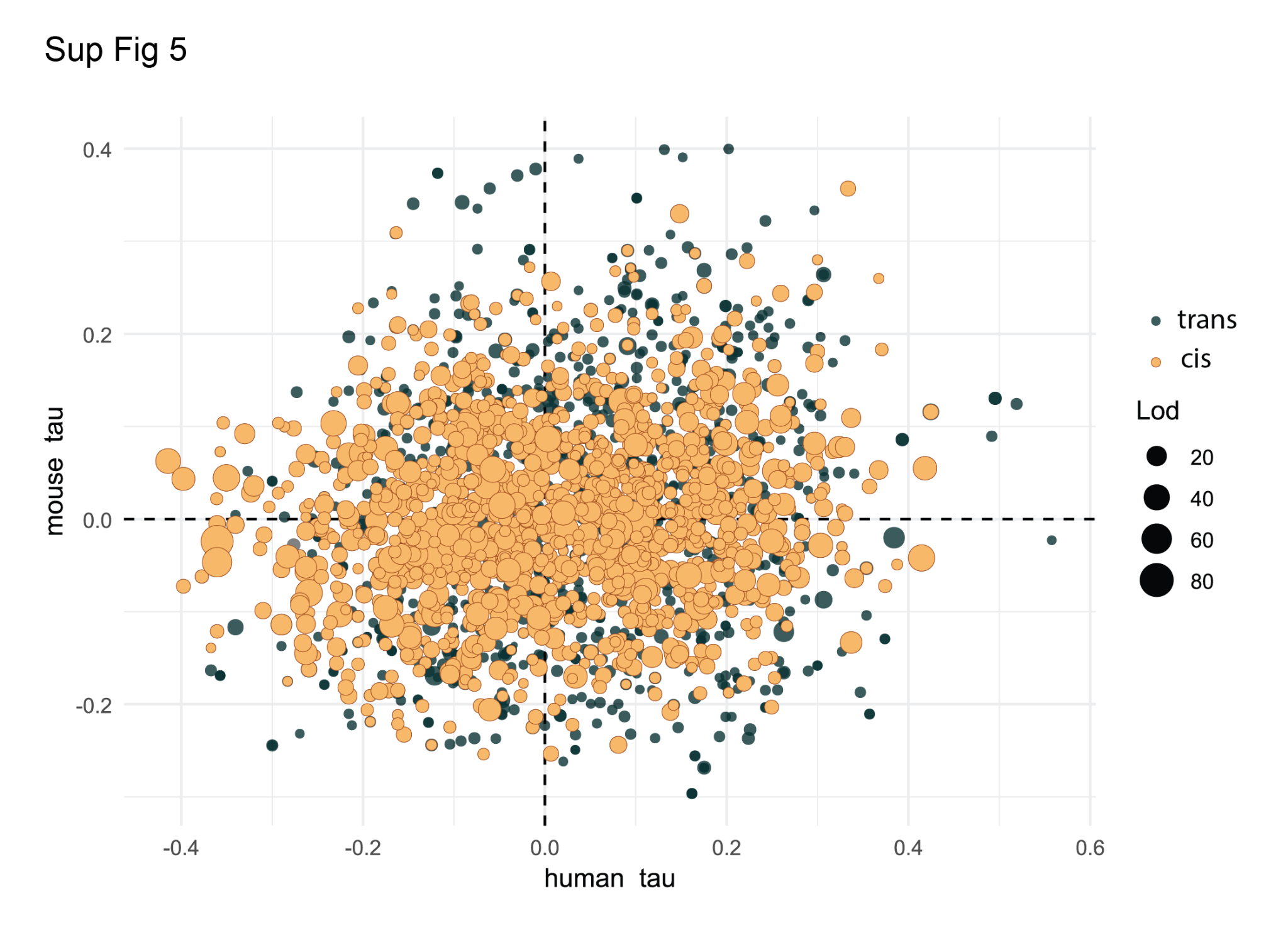

**Sup. Figure 5. Concordance of protein-steatosis correlations between mouse and human liver**

Scatter plot comparing Kendall’s tau correlation (with steatosis/liver fat) calculated in the human cohort (x-axis) versus the mouse cohort (y-axis) for orthologous proteins. Points are colored by pQTL type (cis, yellow; trans, green) and sized by LOD score of the corresponding mouse pQTL. A substantial fraction of orthologs fall in the upper-right and lower-left quadrants, indicating sign-concordant correlation with steatosis across species.

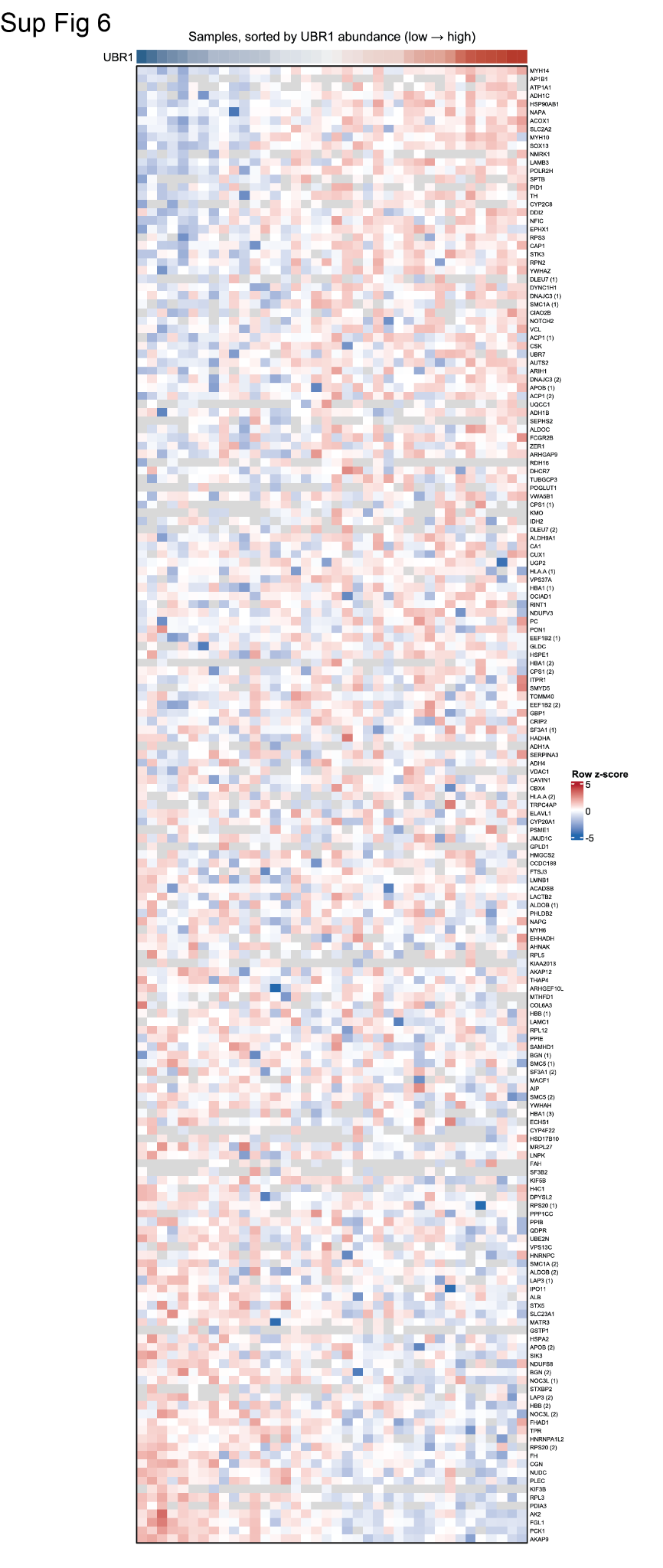

**Sup. Figure 6. Ubiquitination site abundance across human liver subjects.**

Heatmap of ubiquitination site abundance (row z-score) for all detected sites, with samples (columns) sorted by increasing UBR1 abundance (color bar, top) and rows labeled by gene name; numbers in parentheses distinguish multiple sites detected on the same protein. This expanded view extends the subset shown in Figure 9 to the complete set of ubiquitination sites identified in the human cohort.
